# High-speed TIRF and 2D super-resolution structured illumination microscopy with large field of view based on fiber optic components

**DOI:** 10.1101/2023.05.11.540319

**Authors:** Henning Ortkrass, Jasmin Schürstedt, Gerd Wiebusch, Karolina Szafranska, Peter Mccourt, Thomas Huser

## Abstract

Super-resolved structured illumination microscopy (SR-SIM) is among the most flexible, fast, and least perturbing fluorescence microscopy techniques capable of surpassing the optical diffraction limit. Current custom-built instruments are easily able to deliver two-fold resolution enhancement at video-rate frame rates, but the cost of the instruments is still relatively high, and the physical size of the instruments based on the implementation of their optics is still rather large. Here, we present our latest results towards realizing a new generation of compact, cost-efficient, and high-speed SR-SIM instruments. Tight integration of the fiber-based structured illumination microscope capable of multi-color 2D- and TIRF-SIM imaging, allows us to demonstrate SR-SIM with a field of view of up to 150 × 150 μm^2^ and imaging rates of up to 44 Hz while maintaining highest spatiotemporal resolution of less than 100 nm. We discuss the overall integration of optics, electronics, and software that allowed us to achieve this, and then present the fiberSIM imaging capabilities by visualizing the intracellular structure of rat liver sinusoidal endothelial cells, in particular by resolving the structure of their trans-cellular nanopores called fenestrations.

## 1. Introduction

Super-resolution structured illumination microscopy (SR-SIM) is a fluorescence microscopy technique that attains a spatial resolution beyond the diffraction limit together with a high imaging speed. Its efficient, low-power fluorescence excitation, combined with high fluorescence collection efficiency makes SR-SIM particularly well suited for live cell imaging [1]. SR-SIM doubles the spatial resolution laterally in a 2-beam / 2D-SIM implementation and three-dimensionally in the 3-beam / 3D-SIM configuration, which is already in widespread use in commercial instruments based on the original Gustafsson-Sedat system design [2]. In contrast to other super-resolution microscopy methods, such as stimulated emission depletion (STED) [3] or single molecule localization methods SR-SIM benefits from a lower phototoxicity and allows for imaging speeds with super-resolution that can exceed video-rate [4]. In linear SR-SIM the sample is excited with a 2D or 3D modulated sinusoidal interference pattern with a periodicity near the optical resolution limit.

Current implementations of Gustafsson-style SR-SIM usually generate the interference pattern by utilizing the +/-1^st^ (and, in the case of 3D-SIM, zeroth) diffraction orders of laser beams diffracted by a transmission phase grating or by reflection off a diffraction pattern generated by a spatial light modulator (SLM). The main benefit of using a SLM is that it allows for very rapid changes of the illumination pattern compared to the slow mechanical changes that can be realized with a diffraction grating. This, however, imposes limits to the maximum laser power density that such a system can utilize, because the illumination beam diameter is confined to the size of the active surface of the SLM. Furthermore, this typically limits the angles at which the diffraction orders can be collected and diffraction to higher diffraction orders reduces the laser power available to excite fluorescence. In combination, these circumstances ultimately affect the field-of-view (FOV) where fluorescence is excited in the sample and most realizations of SR-SIM typically have a FOV of ∼40 μm × 40 μm. Alternative approaches to overcome at least some of these limitations rely on the splitting of the two or three illumination beams by separate optical elements. This can, for example, be accomplished by a beam splitter and small MEMS mirrors [5], a Michelson interferometer [6], or a Michelson-type interferometer in combination with hexagonal fiber arrays [7,8].

The approach based on a hexagonal fiber array has several advantages, including 3D-SIM compatibility, since a third beam for the center position can easily be added and it also allows for multi-color SIM operation if broadband single mode fibers are used. The hexagonal fiber array pre-fixes the illumination angles required for SR-SIM by holding pairs of optical fibers with 60° angles between them and the laser beams from each fiber are set to be parallel to the optical axis. In this case the FOV is determined by the numerical aperture of the lenses used to focus the beams to the back focal plane (BFP) of the objective lens, such that the full FOV of the objective lens can be utilized, while other SIM methods providing a large FOV, based on a grating [9] or waveguides [10,11] are rather slow or costly to manufacture. In addition, the use of non-diffracting optics enables one to retain much higher laser powers where losses are only limited by the fiber coupling efficiency. The flexibility provided by the separate beam alignment optics furthermore allows for optimal alignment of the interference pattern in the sample plane to achieve high modulation amplitudes. This permits optimal extraction of the maximum amount of high frequency information in the image reconstruction. This flexibility also simplifies total internal reflection fluorescence excitation (TIRF)-SIM with a maximum resolution enhancement. So far, however, current fiber-based implementations are limited by the FOV and phase stability [7,8].

Here, we demonstrate the combination of all the advantages of fiber-based SR-SIM illumination, resulting in an over 10× larger FOV for coherent SIM, fast transition times between angle and phase changes of the illumination pattern, multicolor imaging capability, and quick extension to TIRF-SIM, providing <100 nm spatial resolution. The fiber optic system is readily compatible with different objective lenses, since the hexagonal fiber array can be reproducibly adjusted to adapt for different BFP positions. We demonstrate the performance of this system by imaging the nanoscale morphology of primary liver sinusoidal endothelial cells (LSECs) in 2D-SIM and TIRF-SIM mode.

## 2. Results and Discussion

### 2.1 Optical system

The fiberSIM setup is powered by three lasers (491 nm, 100 mW, Cobolt Calypso 100; 561 nm, 200 mW, Photontec Berlin) and a 639 nm, 300 mW, Photontec Berlin), which are combined with dichroic filters (LaserMUX, Semrock), and passed through an acousto-optic tunable filter (AOTF, AOTFnC-VIS-TN, AAOptoelectronic) to acousto-optically select wavelengths and adjust the laser power. The beam is then coupled into a high-power polarization maintaining fiber (PM fiber, OZ Optics) with coupling efficiencies of >50% for each wavelength by an achromatic fiber-coupler (60SMS-1-4-M7.5-01, Schäfter & Kirchhoff). The PM fiber is connected to a custom-built fiber switch (see Fig. 1), where the beam is collimated and split twice by non-polarizing beam splitter cubes (30/70 and 50/50, Thorlabs Inc.). The first reflected beam with 30% of the optical power is coupled to a fiber that is used as a center beam to align the optical setup between the hexagonal holder and the objective lens and that can be used for 3D-SIM. The two beams with 35% of the optical power, each, are coupled to one of three pairs of PM fibers that are used as side beams to generate the interference pattern in the sample. The corresponding fiber pair for each illumination angle is selected by two galvanometric mirrors (QS7X-AG, Thorlabs Inc.). The scan angle of 1.5° optically was kept minimal to achieve high switching speeds. One side beam is modulated by a custom-built, MEMS-based phase shifter (discussed in detail below) in its optical path length to shift the interference pattern in the sample. The MEMS mirror was driven electrically such that it performs z-shifts instead of angular tilts for this purpose (see below). All beams are coupled by achromatic three-axis fiber couplers (60SMS-1-4-M7.5-01, Schäfter & Kirchhoff) to broadband, high power polarization maintaining single mode fibers (OZ Optics). The beams are outcoupled, collimated and individually focused by a custom-built hexagonal holder which defines the separation between the individual fibers of each pair and fixes the angle of the illumination pattern. Each fiber connector in the outcoupling hexagonal holder is adjusted individually (rotated) to orient the polarization of the beams azimuthally (resulting in s-polarized beams at the sample plane). Both lenses for each beam path (40 mm and 100 mm focal length, Edmund Optics) are laterally adjustable with a custom-built opto-mechanic. This allows precise alignment of the beam position and incidence angle in the back focal plane (BFP) of the objective lens, which is especially important for TIRF-SIM. The second lens in each beam path can be axially shifted to precisely align the axial position of the beam focus in the BFP. This allows to quickly adjust for different objective lens brands with different BFP positions without changing the alignment of the setup, which is a feature that is currently missing from most other SR-SIM implementations and restricts them to the use of a small number of objective lenses. The beam foci are projected via a telescope into the BFP of the objective lens. An iris aperture in the conjugated image plane allows to crop the illumination FOV to the size of the image sensor. The excitation beams are reflected upwards into the objective lens of a custom-built inverted microscope by a quad band polarization maintaining dichroic filter (F73-420PH, AHF AG). The objective lens is either chosen for a large FOV as a 40× 1.4NA (UPLXAPO40XO, Olympus) or for TIRF-SIM as a 60× 1.5NA (UPLAPO60XOHR, Olympus). The fluorescence signal is collected by the same objective lens, transmitted by the dichroic mirror, filtered with a quad band emission filter (F72-891, AHF AG) and imaged onto the sCMOS camera sensor (pco.edge 4.2 CLHS, PCO AG) via a tube lens. With the 40× objective lens we use an achromatic tube lens with 400 mm focal length (Qioptiq) to fulfill the Nyquist criterion and with the 60× objective lens a Ploessl-type lens configuration composed of two achromatic lenses with 500 mm focal length (Thorlabs Inc., 252 mm effective focal length), each. The projected pixel size is 77.3nm for the 60x objective lens and 73.1 nm for the 40x objective lens. The FOV is currently not limited by the excitation beam diameter but the detection path. With the 40x objective lens the full sensor (150 μm projected size) is used, with the 60x the FOV is limited by the planarity of the objective lens to 100 μm. The system is electronically controlled by an ATmega328P microcontroller that triggers the camera and sets an AD5628 digital-to-analog converter (DAC, Analog Devices), which provides analog voltages for controlling the galvo mirrors, the MEMS phase shifter and the AOTF with update rates of up to 25 kHz. The microcontroller allows precise control of the electro-mechanical components with sub millisecond time resolution. The microcontroller communicates with a custom-written Python program on the main computer controlling the instrument, which offers a graphical user interface and integrated control of the piezo stage and the microcontroller hosted functions. The camera is read via MicroManager software [12,13].

**Fig. 1.**
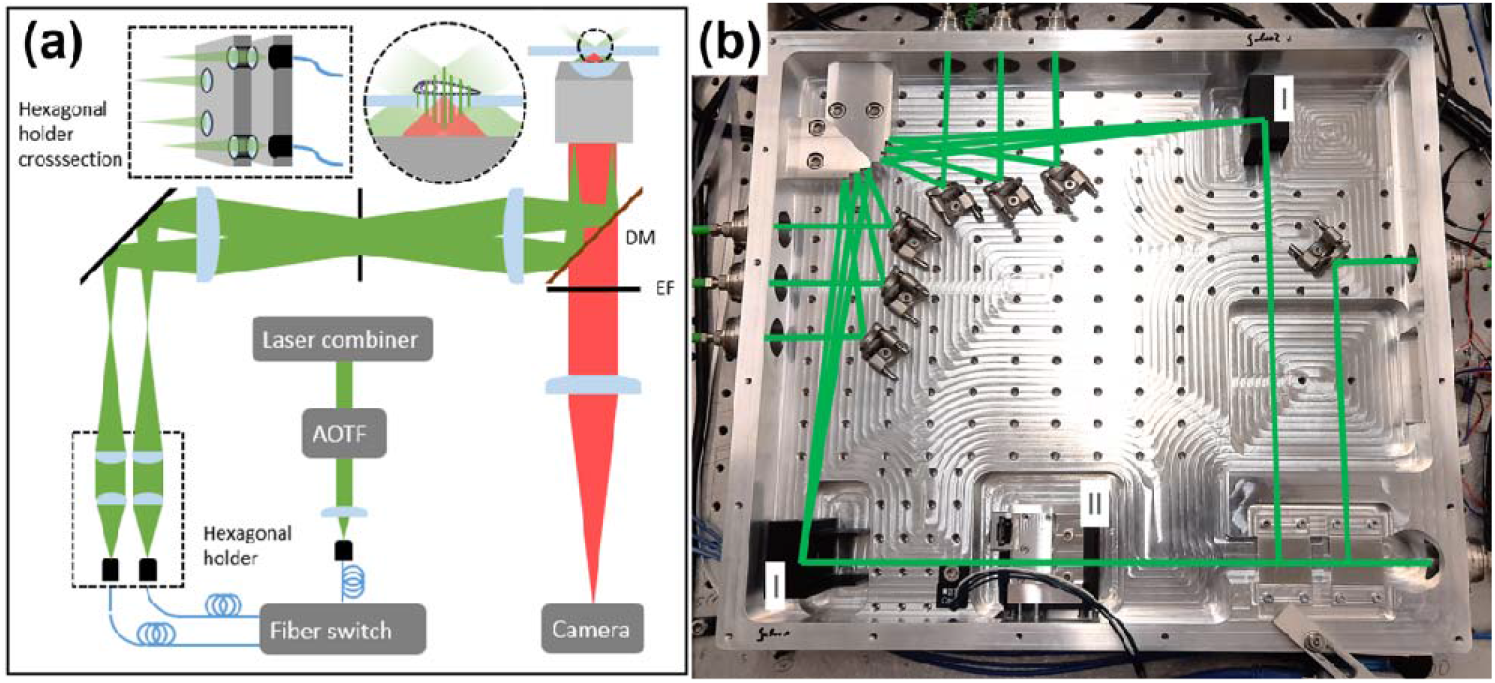
(a) Schematics of the fiberSIM system. A set of lasers (491 nm, 561 nm and 639 nm) is combined and filtered by an AOTF and fiber-coupled to the custom-built fiber switch. (b) Photograph of the fiber switch. In the fiber switch, the free-space laser beam is split to two fibers for the side beams. The corresponding fiber pair for one of the 3 illumination angles is selected by galvanometric mirrors (I). One side beam path is modulated in its optical path length with a MEMS-based phase shifter (II). The beams exiting the fiber pairs are collimated and focused by a hexagonal holder which readily arranges the fiber pairs at 1 of 3 different angles for angle selection. The beam foci are projected in the back-focal-plane of the objective lens by a relay telescope via a polarization maintaining dichroic filter. The fluorescence is epi-detected, filtered and focused by a tube lens onto the image sensor.

### 2.2 Selection of the broadband polarization maintaining optical fibers

The PM fibers are chosen to exhibit a high extinction ratio of >18 dB in order to enable a high modulation depth for the SR-SIM interference pattern. In order for these fibers to be used with all laser wavelengths, we selected broadband fibers that allow for this extinction ratio for a single mode for the entire wavelength range from 488 nm to 650 nm. The fiber end face through which laser light is coupled into the fiber enables coupling laser powers of up to 300 mW. We experienced solarization of fibers that do not have a pure silica core, but a doped core, resulting in a decreased transmission and extinction ratio. This manifests itself as a drop in the extinction ration after several hours of exposure to 491 nm light. This effect was not noticed with pure silica core fibers. Since the fibers have a length of 2 m and the optical path length is dependent on the stress sensitive refractive index of the fiber core and dependent on thermal expansion, it is prone to drift, which appears as a phase drift in the excitation pattern. This drift is typically below 10°/s when the fibers are covered and laid on an optical table. The length of different fibers varies within a few millimeters, therefore the fiber lengths for each pair are matched to be as similar as possible. The coherence length of the lasers, however, needs to be longer than the maximum fiber length difference, therefore single mode DPSS lasers are required.

### 2.3 Phase shifting

Phase stability is among the most crucial parameters for successful SR-SIM microscopy. If the absolute phase drifts within the time that is required to acquire all raw images for SR-SIM reconstruction, then the resulting reconstructed image is prone to exhibit reconstruction artifacts. For this reason we investigated a number of ways by which phase shifting can be accomplished and settled on a microelectromechanical systems (MEMS)-actuated mirror (Mirrorcle Tech. Inc.) with 1.2 mm diameter, which provides excellent phase stability at an acceptable cost. The mirror is placed in a CNC-machined aluminum housing (27x45x18 mm^3^) as shown in Fig. 2a. In this arrangement the incoming beam is first deflected by 84° by a right-angle prism mirror onto the MEMS-mirror. After reflection on the MEMS mirror it is returned onto the same axis as the incoming beam with another right-angle prism mirror. The low angle of incidence of 6° on the MEMS-mirror causes a lateral beam displacement of <50 nm for phase shifts, which is negligible for the fiber-coupling afterwards.

**Fig. 2.**
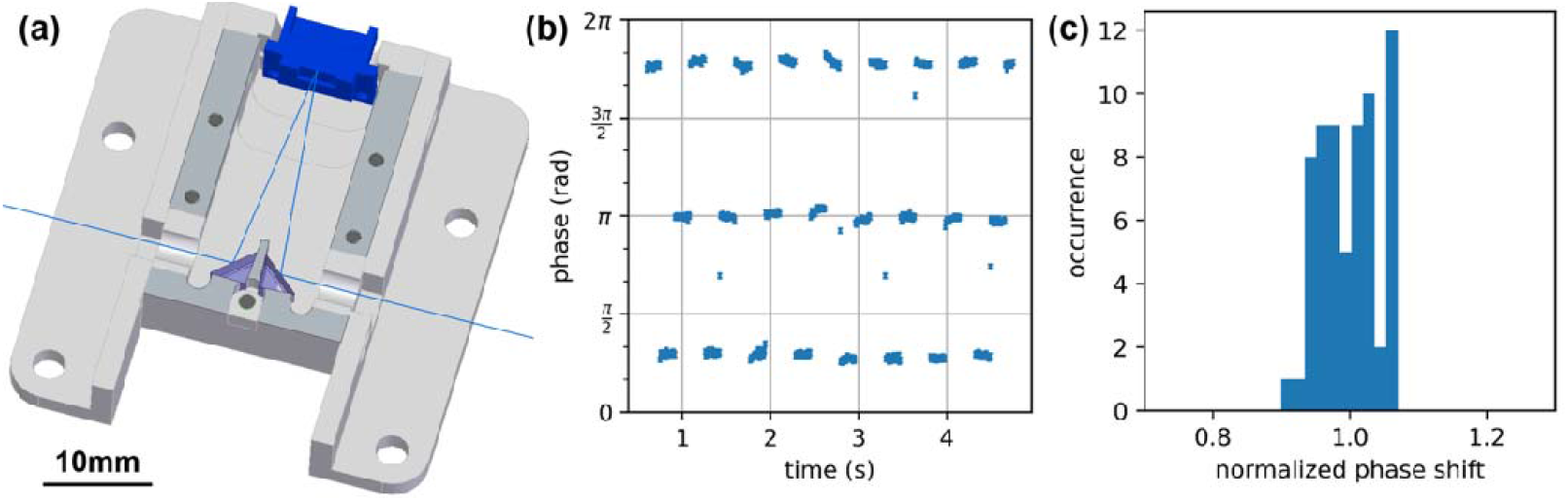
Characterization of the custom-built phase shifter. (a) Cross section of the 3D-CAD model of the MEMS-based phase shifter. The incoming beam is reflected onto the MEMS mirror (assembly shown in blue), which is actuated in axial direction. The reflected beam is directed back onto the same axis as the incoming beam. (b) Measurement of the reproducibility of the relative phase of the excitation pattern as shifted by the phase shifter over time. (c) Histogram of the relative phase shift, indicating that it displays a standard deviation of 4%.

The MEMS mirror is typically used to provide a gimbal-less bidirectional tilt for two perpendicular axes, therefore it has two actuators per axis. It is controlled such that only the actuators of the x-axis are used and one of the two actuators is inverted, to not tilt, but axially displace the mirror. The MEMS actuator is supposed to be controlled with an 80 V offset voltage. In the manipulated configuration, the voltage offset causes a slow angular drift of the mirror (>1 mrad), therefore the actuators were controlled by a voltage of 0-10 V for inducing the phase shift. As can be seen in Figs. 2b and 2c the repeatability of the phase shift is 96% in this low voltage operation mode compared to 99% in the high voltage operation mode. Because the resonance frequency of the mirror is 2.6 kHz, the applied voltage waveform is low pass filtered to avoid ringing of the mirror surface, which could result in intensity ringing of the coupled light.

### 2.4 Switching speed and stability of the fiber switch unit

The fiber switch unit is used to select and switch the interference pattern for the 3 angles used to acquire 2D-SIM images and its stability is critical to maintain a high modulation pattern depth in the sample plane. The speed of the transition between illumination patterns is limited by the switching time of the galvo-mirrors and the MEMS phase shifter. When rotating the galvo-mirror to select an output fiber, the deadtime between fiber coupling is 200 μs and the residual ringing of the coupled intensity is below 10% after 400 μs. This is achieved due to a small scan angle of 1.5° and is equal for both galvo mirrors. The phase shifter is limited by the speed of the axial displacement of the MEMS mirror. The ringing of the phase is below 5% after 1ms. This allows switching speeds of up to 1 kHz while the setup is currently limited by the maximum frame rate of the camera, which is 100 Hz for full frame image acquisition. This can be increased by using two or more cameras for imaging [4]. The reproducibility of the coupled fiber intensity by the galvo-mirror is better than 99%, the reproducibility of the phase shift is above 95%. Since the optical beam path in the fiber switch is up to 800 mm for each beam, the drift stability of all optical components is critical to maintain a constant coupling efficiency of above 50%. Equal coupling efficiencies between beam pairs are needed for a maximum modulation depth. For this reason, the housing of the fiber switch is machined out of a single block of aluminum alloy.

### 2.5 Sample preparation

Cryo-preserved rat LSECs were prepared as described in [14] and stored at -80°C. For thawing and seeding, a vial with LSECs is placed in an incubator at 37°C until nearly all the ice is thawed. The cells are gently pipetted drop-wise to 25 ml of pre-warmed Dulbecco’s Modified Eagle Medium (DMEM) and centrifuged at 50g for 3 minutes to remove any hepatocytes remaining from the cell isolation. The supernatant containing LSECs is used for a second centrifugation step at 300g for 8 minutes. The cell pellet is resuspended in 4 ml - 7 ml DMEM and 1.5 ml (∼ 100.000 cells per cm^2^) of the cell solution is pipetted onto a fibronectin-coated #1.5 glass coverslip. The coverslip surface is coated with fibronectin (0.2 mg/ml) in phosphate buffered saline (PBS) containing 2 mM ethylenediaminetetraacetic acid (EDTA) for 1 hour at room temperature and washed with PBS afterwards. After allowing the cell suspension to incubate on the glass coverslip for 1 hour at 37°C and 5% CO_2_, the coverslip was washed with pre-warmed DMEM and incubated for another 2 hours before fixation with 4% formaldehyde in PBS for 10 minutes at room temperature. The first step of the dual color fluorescent staining of LSECs as shown in Fig. 3 and Fig. 4 is the staining of the plasma cell membrane. The fixed cells were stained with BioTracker 655 Red Cytoplasmic Membrane Dye (SCT108, Sigma) diluted 1:200 in PBS for 1 hour at room temperature. The cells were washed twice in PBS before staining the actin cytoskeleton. The LSECs were incubated in a 1:40 dilution of Phalloidin CF568 (00044-T, Biotium) in PBS for 2 hours at room temperature. After the staining process was completed, the cells were washed 3 times with PBS. The single color actin staining of the LSECs as presented in Fig. 5 is done with a permeabilization step. The cells were incubated in 0.1% Triton X-100 in PBS for 10 minutes at room temperature and then washed 3 times in PBS. Afterwards they were stained in a 1:40 dilution of Alexa Fluor 488 Phalloidin (A12379, ThermoFisher) in PBS.

**Fig. 3.**
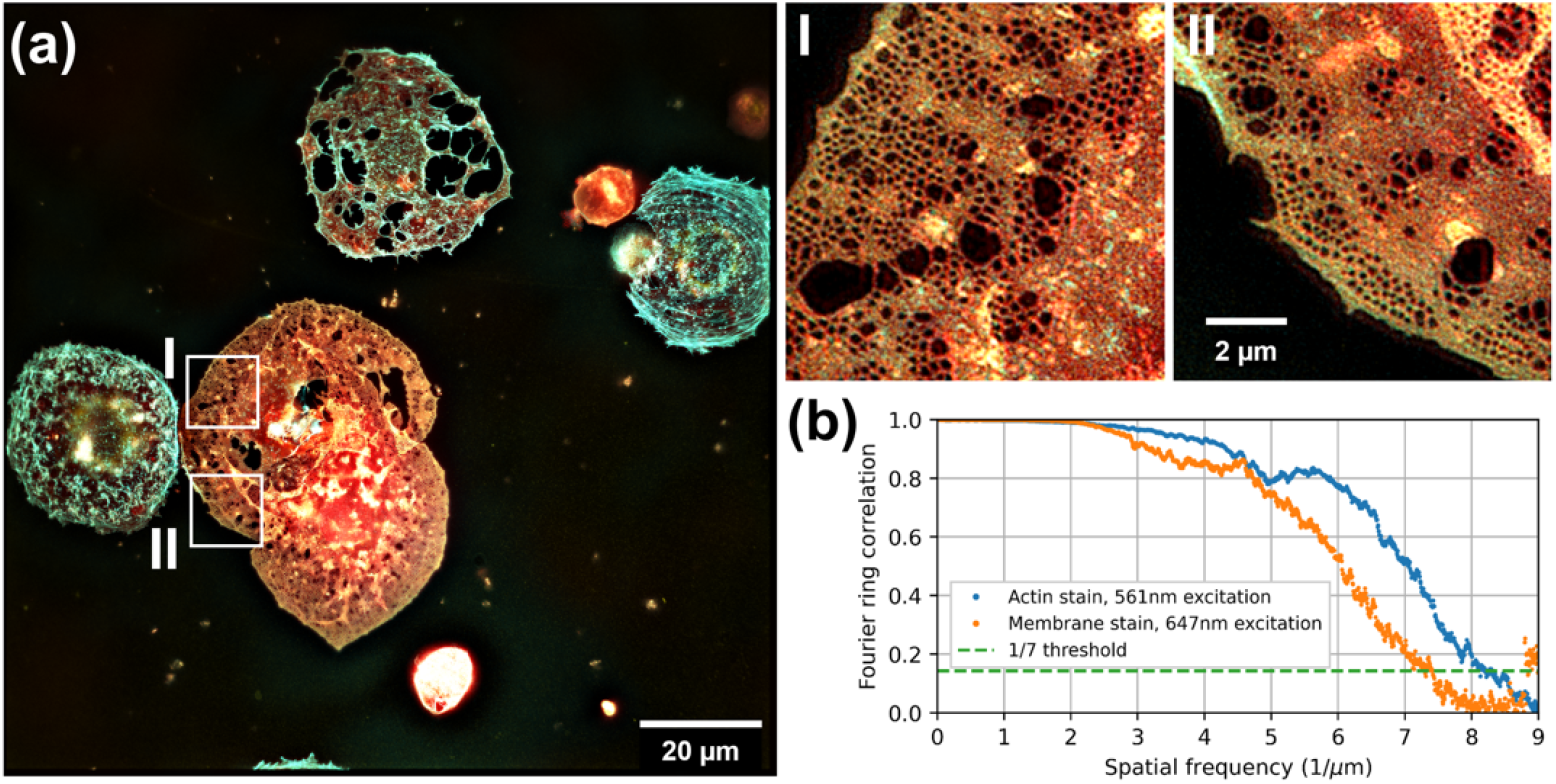
(a) Multicolor 2D-SIM image of fixed LSECs stained with phalloidin CF568 (actin, cyan) and BioTtracker 655 (membrane, red), acquired with a 40× 1.4NA objective lens. The full image data of the sCMOS camera sensor was reconstructed to obtain this image (150 μm×150 μm FOV). (b) The spatial resolution as determined by Fourier ring correlation results in a resolution of 121 nm (red channel) and 137 nm (deep red channel). Both insets (I, II) have the same size.

**Fig. 4.**
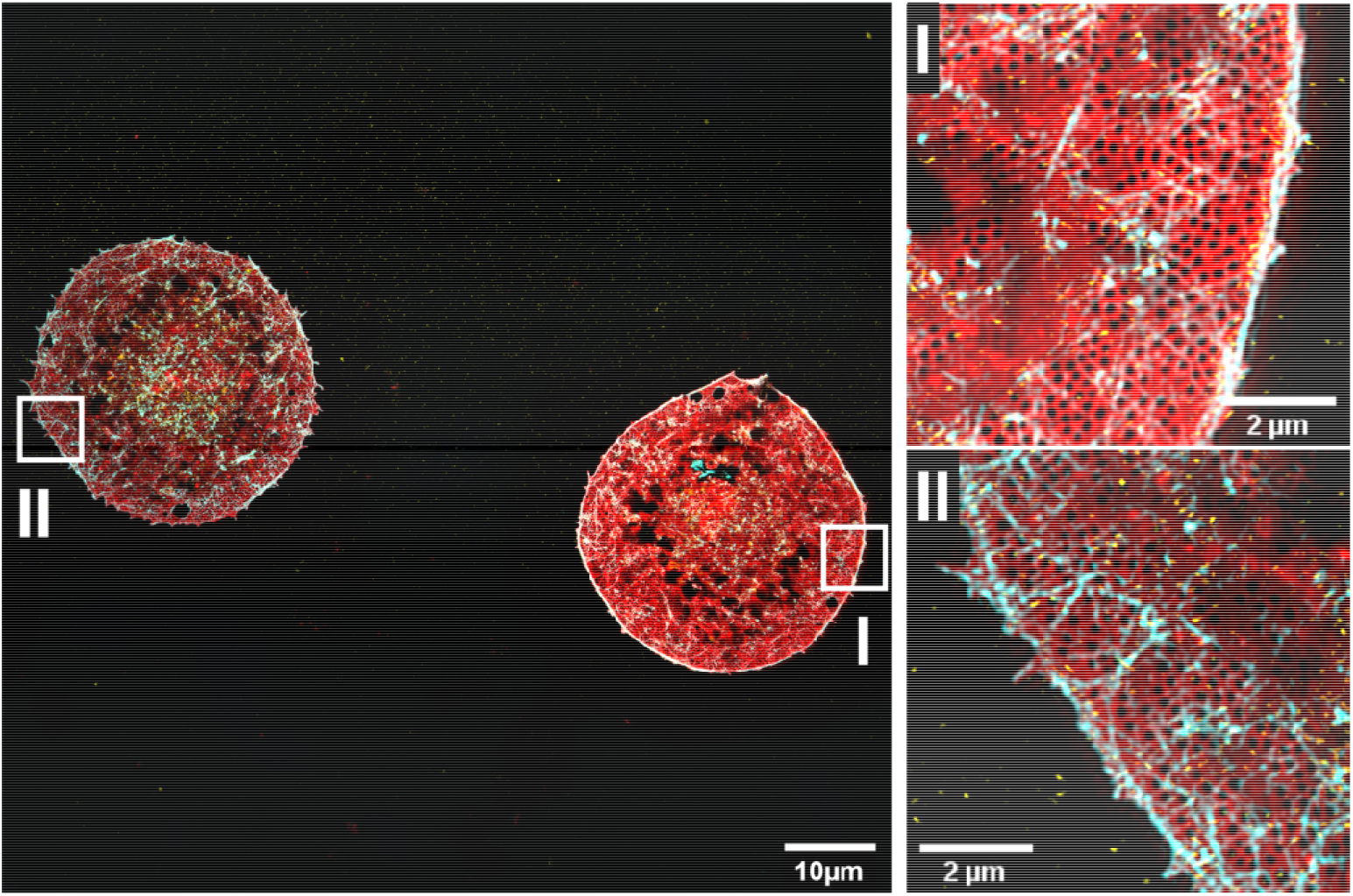
TIRF-SIM image of LSECs seeded and fixed on a glass cover slip. The cells are stained with phalloidin CF568 (actin, cyan) and BioTracker 655 (membrane, red). The TIRF-SIM reconstruction provides superior results for all three wavelengths over the full FOV. The spatial resolution is 140 nm (membrane, red channel), 117 nm (actin, cyan channel) as determined by Fourier ring correlation.

**Fig. 5.**
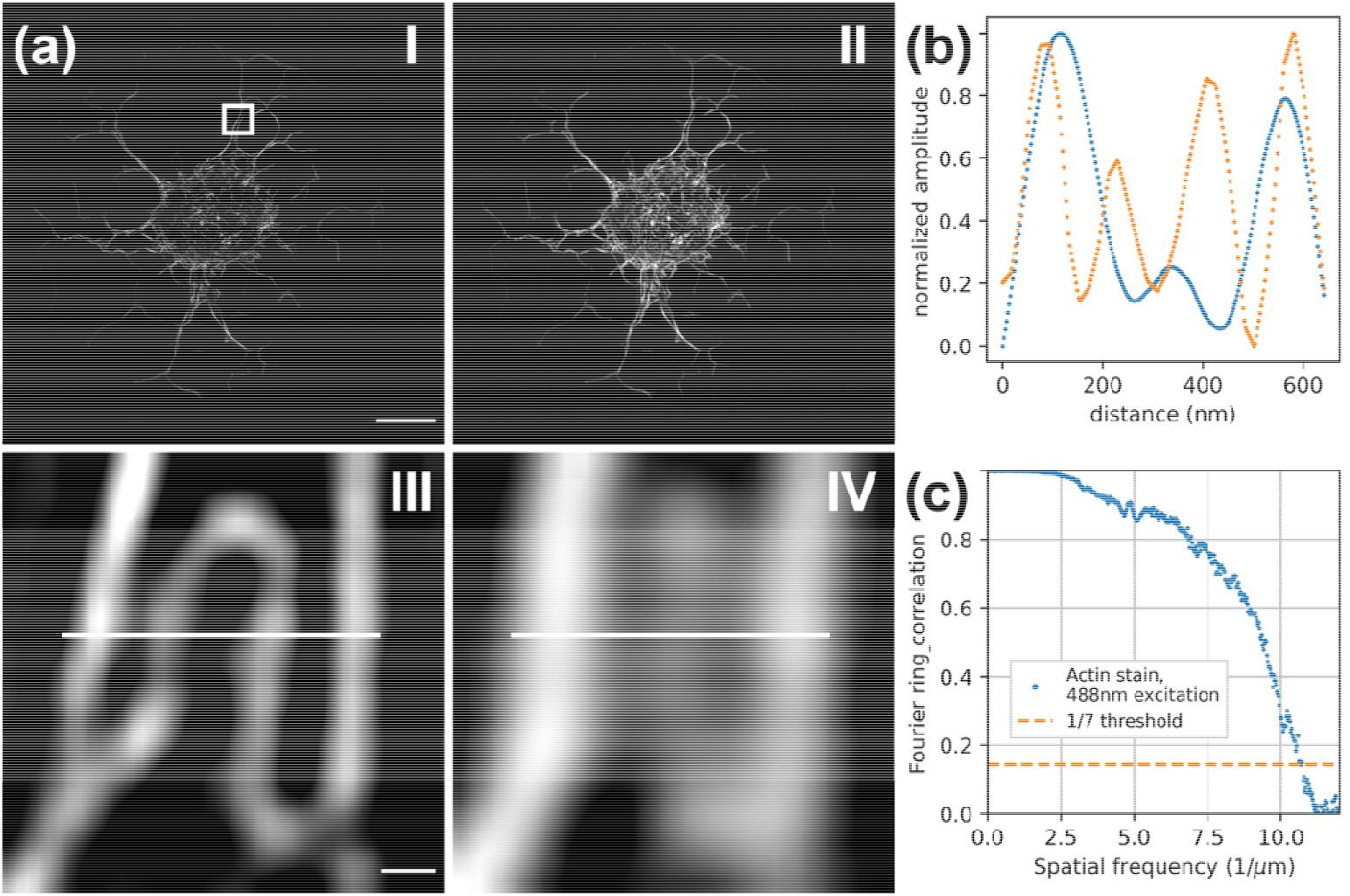
(a) TIRF-SIM (panel I and III) and filtered widefield (panel II and IV) images of a fixed rat LSEC stained with phalloidin AF488 (actin) and imaged with maximum linear SIM resolution improvement. (b) The line plot across the actin filaments shown in panel III exhibits that nearby filaments are well separated and distinguished. (c) The spatial resolution as measured by FRC is 93 nm. The scale bars are 5 μm (panel I, II) and 100 nm (panel III, IV), the FOV is 37 × 37 μm^2^(panel I, II).

### 2.6 SR-SIM imaging results

The fiberSIM setup in its current implementation enables the acquisition of SR-SIM data with a large field of view (FOV). When compared to existing SIM microscopes, this extended FOV is accomplished by the choice of the rather short focal length provided by the second lens inside the hexagonal fiber holder. The performance of the fiberSIM system was demonstrated by imaging fixed LSEC on a glass coverslip surface. SR-SIM raw images were reconstructed using the fairSIM software [15], available as a FIJI plugin. For the reconstruction of the SR-SIM images 3 phase steps per angle are required, resulting in 9 raw images per super-resolved image. The increased FOV in combination with the 40× 1.4NA objective lens allows us to utilize the full sCMOS camera sensor with 2048 pixel × 2048 pixel resulting in a FOV of 150 μm × 150 μm during image acquisition as shown in Fig. 3a. This enables the reconstruction of SR-SIM images without a significant drop in signal intensity across the entire FOV even with multicolor fluorescence excitation (see Fig. 3a). The illumination pattern distortion for this extended FOV is low enough to not affect the reconstruction in 2D-SIM and the resulting images exhibited little to no reconstruction artefacts throughout the entire FOV. Figure 3a shows a multicolor 2D-SIM image of 5 fixed LSECs, where the actin cytoskeleton is stained with phalloidin CF568 (shown in cyan) and the plasma membrane is stained with BioTracker 655 (shown in red). As can be seen from the enlarged insets in Fig. 3, individual fenestrations, i.e. nano-sized trans-cellular holes in the cell body of LSECs, can be resolved with our fiberSIM system, allowing for the analysis of a large number of cells, simultaneously, compared to typical single-cell size FOV in current SR-SIM systems. The fenestrations are arranged in groups of about 5-100, known as sieve plates, as can be seen in both insets. The spatial resolution of the images is 121 nm for the actin channel and 137 nm for the plasma membrane channel as indicated by the Fourier ring correlation (FRC, [16]) results shown in Fig. 3b. The bright spots in inset I are mostly due to agglomerations of the plasma membrane stain. The brighter area in the upper right corner in the image in inset II is part of another LSEC that has adhered to the surface right below the cell highlighted by inset I. Part of this lower cell partially overlaps with the upper cell as can be seen in Fig. 3a. The larger, irregular gaps in the plasma membrane as most evident in the uppermost LSEC in Fig. 3a are often observed in cryopreserved LSECs grown on fibronectin-coated glass surface.

When using a 60x 1.5NA objective lens, the FOV is limited to a size of 100 μm x 100 μm without reconstruction artefacts. This reduced FOV is due to the limited raw image planarity of the objective lens. The modulation depth which we obtained with this objective lens was greater than 70%, enabling high quality SR-SIM image reconstruction. The effective FOV was the same for TIRF- and 2D-SIM images. Since the chromatic aberration of the 60× objective lens is not optimized for the deep red channel, a focus shift of 2 μm must be considered during image acquisition. We do not observe any further aberrations in the deep red channel. Accordingly, Fig. 4 shows a TIRF-SIM image of two LSECs seeded and fixed on a glass cover slip. Here, the cells are stained with phalloidin CF568 (actin, cyan) and BioTtracker 655 (membrane, red). As can be seen from the enlarged insets in Fig. 4, TIRF-SIM reconstruction provides an even better spatial resolution taking advantage of both, the higher numerical aperture of the objective lens and the finer interference pattern in TIRF mode resulting in superior contrast for fenestrations. According to the results of a Fourier ring correlation analysis of these images, the spatial resolution is 140 nm for the deep-red plasma membrane channel and 117 nm for the actin cytoskeleton shown in cyan.

Since TIRF-SIM allows for a theoretical resolution improvement of more than 2x and the 1.5NA objective lens allows a high angle of incidence of the excitation beams, we showed that the resolution limit can be pushed to 93 nm for the green fluorescence channel. With an excitation wavelength of 491 nm and the beam foci placed at the edge of the BFP (corresponding to an angle of incidence of 81° with the 1.5NA objective lens), we expect a pattern spacing of 163 nm in the sample plane. This results in an effective resolution improvement of 2.07, while 2.08 is the physical limit given by the excitation and emission spectrum. This fits well with the actual pattern spacing of 163.2 nm and the resolution improvement of 2.07 as measured in Fig. 5. Fig. 5a shows the phalloidin AF488 labeled cytoskeleton of a fixed LSEC in TIRF-SIM mode (I) and, for comparison as filtered widefield image (II). The enlarged insets shown in III and IV demonstrate the greater than 2x resolution improvement, which reveal that the actin cytoskeleton splits and forms a loop in between the two filaments on the left and right hand side of the image, which provides information that cannot be obtained from the widefield image. The associated cross sections are shown in Fig. 5b and clearly reveal four well isolated actin fibers in the TIRF-SIM channel, whereas an apparent three fibers can barely be identified in the widefield cross section. Here, the resolution limit is 93 nm as determined by Fourier ring correlation.

## 3. Conclusions

We have presented an innovative design of a 2D- and TIRF-SIM microscope offering a significantly increased FOV, high modulation pattern contrast, and a spatial resolution below 100 nm. The system is based on an interferometer housed in a rugged aluminum unibody, where laser beams throughout the visible spectrum can be split and sent to different fiber couplers by using fast-switching galvo mirrors. The phase of the SIM pattern is controlled by a compact MEMS-based phase shifter, providing excellent long-term stability and repeatability. Broadband polarization-maintaining optical fibers guide the laser light to a freely positionable hexagonal fiber holder, where the laser light is converted to free space beams with short focal lengths. The entire system is modular (consisting of 3 base modules: a laser combiner, a fiber switch unit, and a compact inverted microscope) and, due to its efficient use of fiber optics, enables the versatile and free arrangement of the individual units resulting in a SR-SIM system with a small overall footprint. The system readily provides multicolor and high-speed imaging capability, which is only limited by the readout time of the sCMOS camera (100 Hz frame rate for full sensor image transfer, up to 1 kHz for smaller regions of interest). The frame rate could be further increased by using additional cameras. We have demonstrated the performance of this SR-SIM microscope by imaging fluorescently stained primary rat LSECs in 2D- and TIRF-SIM mode, revealing the transcellular holes with diameters around 100 nm, which are characteristic for these cells.

## Acknowledgements

The authors would like to thank Jochen Linnenbrügger for the CAD design of the fiber switch housing and the hexagonal holder. We also thank Dr. Peter Tinning and Dr. Ralf Bauer, both at the University of Strathclyde, for helpful discussions about customizing the control of MEMS mirrors.

